# Optimization of RNA Pepper sensors for the detection of arbitrary RNA targets

**DOI:** 10.1101/2023.06.01.543282

**Authors:** Anli Tang, Anna Afasizheva, Clara Cano, Kathrin Plath, Douglas Black, Elisa Franco

## Abstract

The development of fluorescent light up RNA aptamers (FLAPs) has paved the way for the creation of sensors to track RNA in live cells. A major challenge with FLAP sensors is their brightness and their limited signal-to-background ratio both *in vivo* and *in vitro*. To address this, we develop sensors using the Pepper aptamer, which exhibits superior brightness and photostability when compared to other FLAPs. The sensors are designed to fold into a low fluorescence conformation, and to switch to a high fluorescence conformation through toehold or loop-mediated interactions with their RNA target. Our sensors detect RNA targets as short as 20 nucleotides in length with a wide dynamic range over 300-fold *in vitro*, and we describe strategies for optimizing the sensor’s performance for any given RNA targets. To demonstrate the versatility of our design approach, we generate Pepper sensors for a range of specific, biologically relevant RNA sequences. Our design and optimization strategies are portable to other FLAPs, and offer a promising foundation for future development of RNA sensors with high specificity and sensitivity for detecting RNA biomarkers with multiple applications.

## 1. INTRODUCTION

Fluorogenic RNA aptamers have emerged as a powerful tool for real-time monitoring of RNA expression, localization, and dynamics in living cells. These RNA molecules, also known as fluorescent light-up aptamers (FLAPs), are selected to bind to fluorophores and undergo conformational changes that lead to fluorescence enhancement (1). A multitude of FLAPs and cognate fluorophores has been demonstrated to work *in vitro* and within living cells: examples include aptamers known as Malachite Green (2, 3), GFP-mimics Spinach, Broccoli, and Corn (4– 7), Mango I-IV (8, 9), and Pepper (10).

Recent advances in nucleic acid nanotechnology have made it possible to demonstrate FLAPs working as sensors that fluoresce specifically upon hybridization to the intended RNA target (11, 12). These target-responsive FLAP-based RNA sensors are genetically encoded but do not require any modifications to the RNAs of interest (13). FLAP sensors simply need to be transcribed and hybridized to their RNA target, thereby activating their fluorogenic function (sensor ON). To be effective to detect and track RNA molecules in live cell imaging, FLAP sensors should exhibit a low background fluorescence in the absence of target (sensor OFF). A simple strategy to achieve this is to split RNA FLAPs in two non-fluorescent domains, whose assembly is seeded by the target nucleic acid (11, 12, 14–16). Researchers have successfully demonstrated the recognition of endogenous mRNA in mammalian cells and RNA imaging in E. coli using this approach (11, 12, 16). Another approach is that of designing the FLAP so that it fluoresces only upon binding to its RNA target and undergoing a conformational change, while otherwise remaining in the OFF state. This approach has the advantage of requiring transcription of a unimolecular sensor component with high specificity for its target; further, the FLAP secondary structure is programmable via sequence design (17), and its conformational changes can be optimized to maximize the ON/OFF ratio (18–26). Although existing FLAP-based RNA sensors (Table S1) enable the noninvasive visualization of endogenous mRNAs in living cells, imaging targets at low copy number remains a challenge because it requires both brightness of the ON FLAP sensor and low OFF signal.

A promising candidate to build RNA sensors for low abundance targets is “Pepper” (Fig. 1A1), a 43-nucleotide FLAP that exhibits higher brightness and photostability compared to other established fluorogenic aptamers including Broccoli and Corn, and even some fluorescent proteins such as mCherry (10). An “inert Pepper” (iPepper) sensor was previously shown to detect endogenous RNA *in vivo* in different cell types (27). In this system, the target RNA binds and stabilizes the terminal ends of a misfolded Pepper aptamer, resulting in fluorescence activation. The system’s optimization relied on a tandem array of iPepper (8x), and achieved an ON/OFF ratio of approximately 10-fold *in vitro*. To further improve the sensor ON/OFF ratio, alternative sensor design strategies are needed.

**Figure 1.**
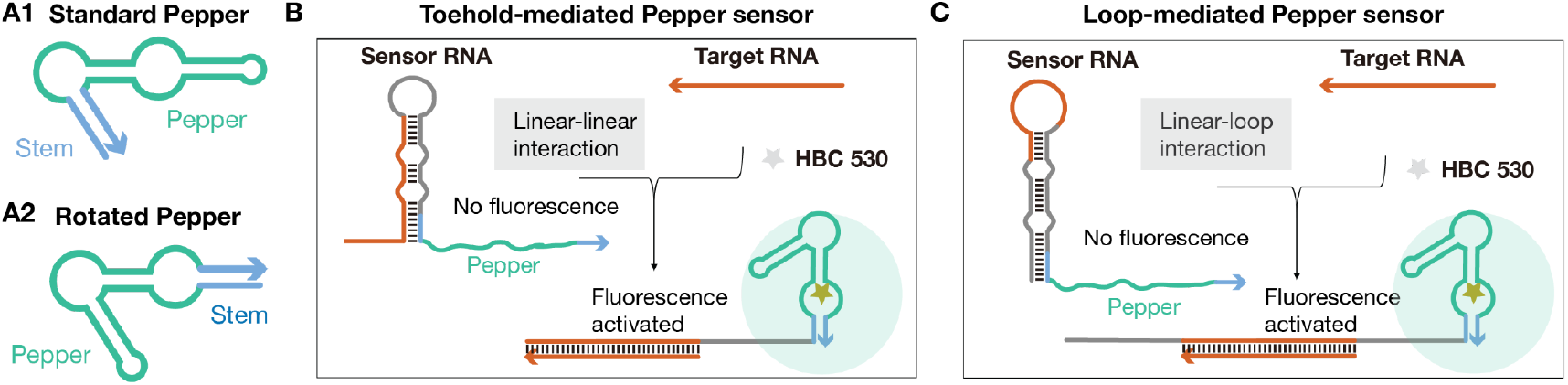
Overview of our Pepper RNA sensor design strategy and its variations. (A1)The secondary structure of the original Pepper aptamer. (A2)The secondary structure of the modified Pepper aptamer. (B)Schematic of the toehold-mediated Pepper sensor. (C) Schematic of the loop-mediated Pepper sensor.

In this study, we introduce and demonstrate design approaches and generalizable optimization strategies for Pepper-based RNA sensors that can achieve up to 300-fold fluorescence enhancement *in vitro*. We demonstrate two types of Pepper sensor designs adapted from previous work demonstrating switchable RNA sensors (26, 28–31), that take inspiration from RNA riboswitches developed for translation control in bacteria (32, 33). In both designs, the 5’ end of the Pepper stabilizing stem is sequestered in the stem-loop structure of the sensor, which destabilizes Pepper and forces it to adopt a non-fluorescent configuration (Fig. 1B and C) (29). The first design, named the toehold-mediated sensor, employs a 5’ toehold domain to initiate the interaction with the target RNA strand, and induce refolding of the aptamer into its fluorogenic conformation (Fig. 1B). The second design, named the loop-mediated sensor, initiates the interaction with the target strand via the loop domain of its hairpin structure (Fig. 1C). Using *in vitro* transcription and plate reader assays, we demonstrate that both sensors can be used to detect target RNA as short as 20-nt with no sequence constraints and a high dynamic range.

Furthermore, we provide strategies to optimize the performance of the sensor for any given target RNA sequences. These strategies aim to reduce the sensor’s leakage in the absence of the target RNA (sensor OFF) and to increase the fluorescence intensity of the sensor when the target RNA is present (sensor ON). Iterative application of these design strategies can result in several hundred-fold increase in the ON/OFF ratio of a sensor. Using these design principles, we generate Pepper sensors for a range of specific, biologically relevant RNA sequences, including a MS2-repeat sequence, Tubulin Alpha 1b (TUBA1B) transcript sequence, metastasis-associated lung adenocarcinoma transcript 1 (MALAT1) sequence, the A-repeat sequence of mouse and human X-inactive specific transcript (Xist/XIST) RNA, and the microRNA miR-302a, miR-294 and miR-124. Finally, we apply our optimization strategies to the Pepper sensor for detecting miR-294, achieving an ON/OFF ratio of approximately 300-fold.

Because our design,characterization and optimization workflow is portable to any other type of FLAP, we expect our results will be immediately useful to build a variety of RNA sensors for detection of biomarkers *in vitro*. The efficacy of our RNA sensors was unaffected when tested in the presence of total RNA from HEK293 cells, suggesting that our design strategies will be applicable to RNA sensing in living cells.

## 1. MATERIAL AND METHODS

### Toehold- and loop-mediated Pepper sensor design

All our Pepper sensor variants were designed with NUPACK (17). We report the scripts in the Supplementary Information file (SI). The sensor strands with the lowest normalized ensemble defects were selected as candidates for *in vitro* experiments.

### Sample preparation

DNA templates for *in vitro* transcription were purchased from Integrated DNA Technologies. DNA oligonucleotides were amplified using Phusion® High-Fidelity PCR Master Mix with HF Buffer (M0531L, New England BioLabs). AmpliScribe™ T7-Flash™ Transcription Kit (ASF3507, Lucigen) was used according to the manufacturer protocol with 4 μL PCR products per 20 μL reaction. RNA was transcribed at 37 °C for 30 min, and the transcription was terminated with the addition of DNase I (Lucigen, ASF3507). RNA targets were purified with Monarch® RNA Cleanup Kit (T2040L, New England BioLabs).

### Pepper sensor screening

Sensors were screened in 96-well assay plates containing 2 μL of sensor RNA, 4 μL of target RNA, 2 μM of HBC 530, 40 mM HEPES, 5 mM MgCl2, 100 mM KCl and water for a final reaction volume of 100 μL. The fluorescence was monitored on a plate reader (BioTek H1) every minute for 2 h at 37 °C. For each sensor variant, the ON signal level was the fluorescence signal measured for the sensor in the presence of the target, whereas the OFF signal level was the fluorescence signal of the sensor alone, in the absence of target. To evaluate the performance of each sensor variant, we report the ON/OFF ratio throughout the paper. The ON and OFF signals were measured at one-minute intervals for two hours, and the ON/OFF were calculated from data at two-hour time points, unless otherwise specified. Although a background control containing all buffer components was measured, it was not subtracted for the ON/OFF calculation. We conducted three replicates of the experiment.

### RNA extraction

Cell-extracted RNA was collected from Hek293 cells. Cells were maintained in media composed of 10% FBS (Life Technologies, 10099141), 100mML-Glutamine (GIBCO, 25030-081), 1x MEM Non-Essential Amino Acids (NEAA) (GIBCO,11140-050), 0.1mM 2-Mercaptoethanol (GIBCO, 21985-023) in DMEM (Sigma, D6429). Cells were harvested, washed with DPBS and collected with Direct-zol (Zymo Research). RNA was extracted using the Direct-Zol RNA miniprep kit (Zymo Research) and quantified using a NanoDrop spectrophotometer ND-1000 (Thermo Scientific).

## 2. RESULTS

### 2.1 Design of toehold-mediated Pepper sensors

The Pepper aptamer includes a fluorogen binding domain for its conjugate dye (HBC 530), which generates the fluorescent response, and a stem domain that stabilizes the aptamer structure (Fig. 1A1). First, we wanted to test whether changes in the stem sequence affect the aptamer fluorescence, so we evaluated three Pepper aptamer variations with mutated stem sequences (Fig. S1A), following the workflow in (27, 29). In addition to the original Pepper aptamer, here referred to as standard Pepper (Fig. 1A1), we considered circular permutations of Pepper, here referred to as rotated Pepper (Fig. 1A2). We expected that the rotated Pepper, previously considered in (27), would have a higher tolerance to the stem sequence change because it has been shown that alterations at the terminal stem-loop of the Pepper aptamer do not affect its fluorescence (10). Our results showed that both standard and rotated Pepper had consistent fluorescence when their stem sequences were scrambled (Fig. S1B). To begin with the sensor design, we first destabilized the Pepper aptamer and kept it in a non-fluorescent conformation by sequestering the 5’ end of the aptamer stem into a large hairpin structure. We then evaluated approaches to design the sensor and its target-binding region, to achieve the release of the aptamer stem and refolding of the aptamer into its fluorescent conformation.

The first design strategy (Fig. 2A), named the toehold-mediated Pepper sensor, contains a 5’ toehold sequence as part of the target-binding region (orange), the core hairpin structure, and the Pepper aptamer sequence at the 3’ end (blue and green), following a strategy proposed in (28– 30). The Pepper stem is initially sequestered with the **b**\* domain of the core hairpin, thereby preventing binding of the fluorogen HBC 530 to the sensor RNA and resulting in a non-fluorescent (OFF) state in the absence of the target RNA. Upon the presence of target RNA, the toehold region of the sensor RNA initiates hybridization with the target, which mediates the strand-displacement reaction necessary for the conformational change of the sensor. This causes the unwinding of the core hairpin, thereby releasing the **b**\* domain, which then hybridizes with the 3’ end **b** domain, leading to correct folding of the Pepper aptamer and resulting in high fluorescence (ON).

**Figure 2.**
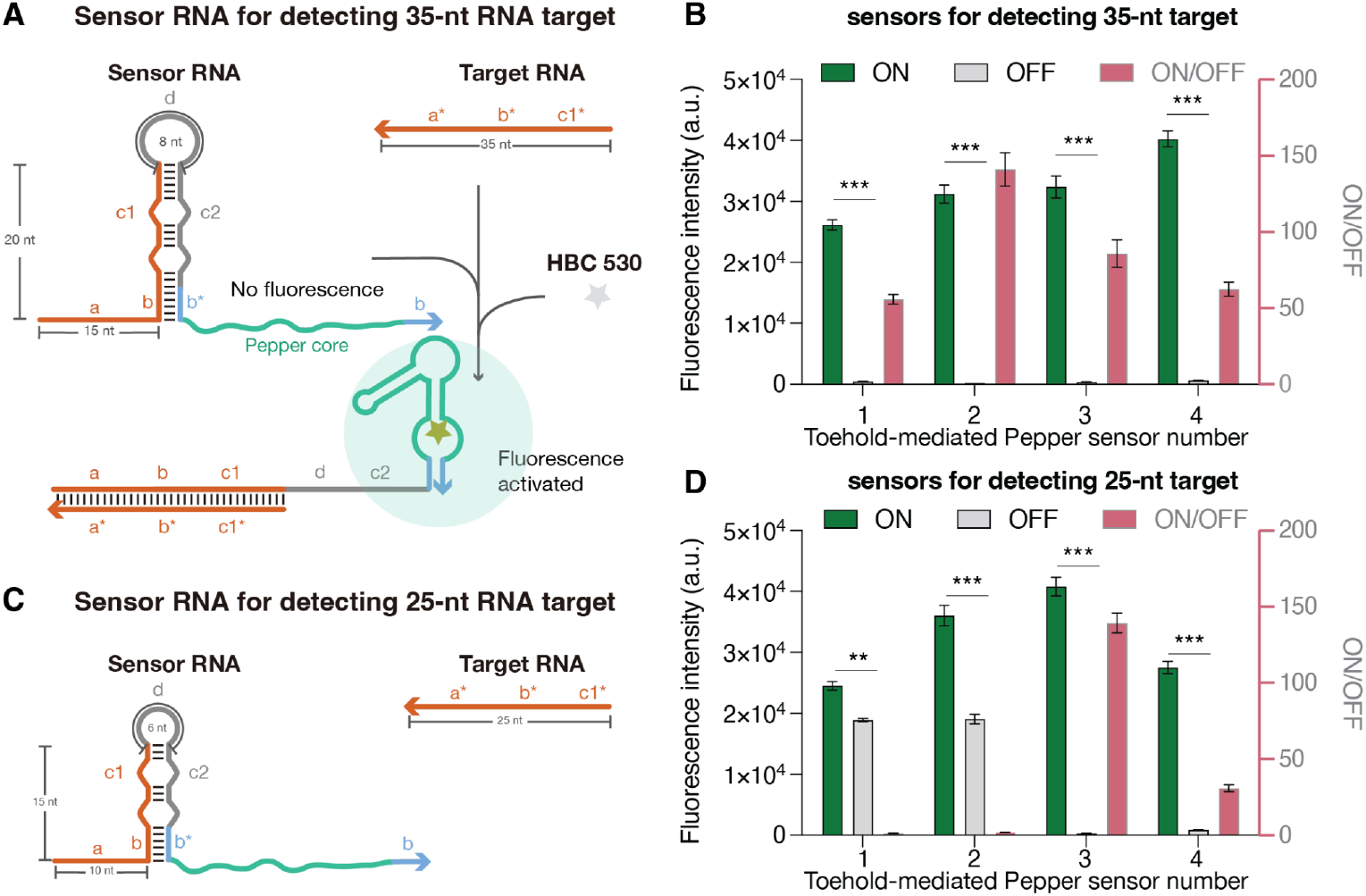
Toehold-mediated Pepper sensors. (A) Detailed schematic of the toehold-mediated Pepper sensor targeting 35-nucleotide target RNA. (B) Best performing toehold-mediated Pepper sensors targeting arbitrary RNA targets. Data represent mean fluorescence intensity from plate reader measurements with sensor alone (OFF) and sensor plus 35-nucleotide target RNA (ON). ON/OFF ratios shown in pink (right y axis). Error bars represent standard deviations from three technical replicates. (C) Schematic of the toehold-mediated Pepper sensor targeting 25-nucleotide target RNA. (D) Best performing toehold-mediated Pepper sensors targeting shorter RNA targets. Data represent mean fluorescence intensity from plate reader measurements with sensor alone (OFF) and sensor plus 25-nucleotide target RNA (ON). Error bars represent standard deviations from three technical replicates. Significance: (***) for p < 0.001, (**) for p < 0.01.

To start, we designed sensors targeting arbitrary RNA sequences because to test the potential of the sensors in ideal conditions when there are no sequence constraints at the toehold, loop (**d** domain) and the **b** domain; the only fixed sequence was the Pepper core highlighted in green (in other words, the target sequence was included in the sequence optimization step). We generated toehold-mediated sensor candidates for detecting 35-nt RNA targets using the NUPACK nucleic acid sequence design package (17) and selected the top four sensor variants with the least ensemble defects from the NUPACK prediction for our *in vitro* plate reader assay (Fig. 2B). We found sensors adopting the rotated Pepper aptamer to be better performing, based on their low OFF signal and high ON signal when activated by their cognate target RNA (Figure 2B), when compared to the ones adopting the standard Pepper aptamer (Figure S1C). Therefore, we employed the rotated Pepper aptamer in subsequent sensor designs.

Next, we developed sensors that could detect shorter RNA targets: small non-coding RNAs, like small interfering RNA (siRNA) and microRNA (miRNA), are 19-to 25-nt in length and play important roles in gene regulation (34). To make it possible to detect RNA targets of comparable length, we designed toehold-mediated sensors with a shorter sensing domain by changing 1) the length of the toehold region from 15-to 10-nt; 2) the length of the stem of the core hairpin from 20-to 15-nt; 3) the size of the loop of the core hairpin from 8-to 6-nt (Figure 2C). We then generated and selected four toehold-mediated sensor variants with shorter sensing domain, and evaluated them through our *in vitro* plate reader assay (Fig. 2D and Fig. S1D). Among the four variants tested, sensor #3 and #4 exhibited better performance with negligible fluorescence leakage in their OFF state when compared with the background signal (Fig. 2D and Fig. S1D). Our results suggest that the impact of sequence variation on sensor performance is greater in the sensor designed to detect 25-nt targets than in the sensor designed for 35-nt targets, possibly because of their shorter sensor domain.

### 2.2 Forward engineering of the toehold-mediated Pepper sensor

Next, we asked whether we could systematically engineer sensors to improve their ON/OFF ratio, and we focused on the variant with the worst performance (Fig. 2D, toehold-mediated sensor #1 for detecting 25-nt RNA). We began by trying to minimize the signal leakage and made changes that include 1) removing the bottom bulge, 2) removing both bulges, and 3) removing the bottom bulge and extending the stem length (Figure 3A). The sensor with a single bulge showed the most notable improvements in its ON/OFF ratio (Figure 3B), and we kept this modification for our following designs.

**Figure 3.**
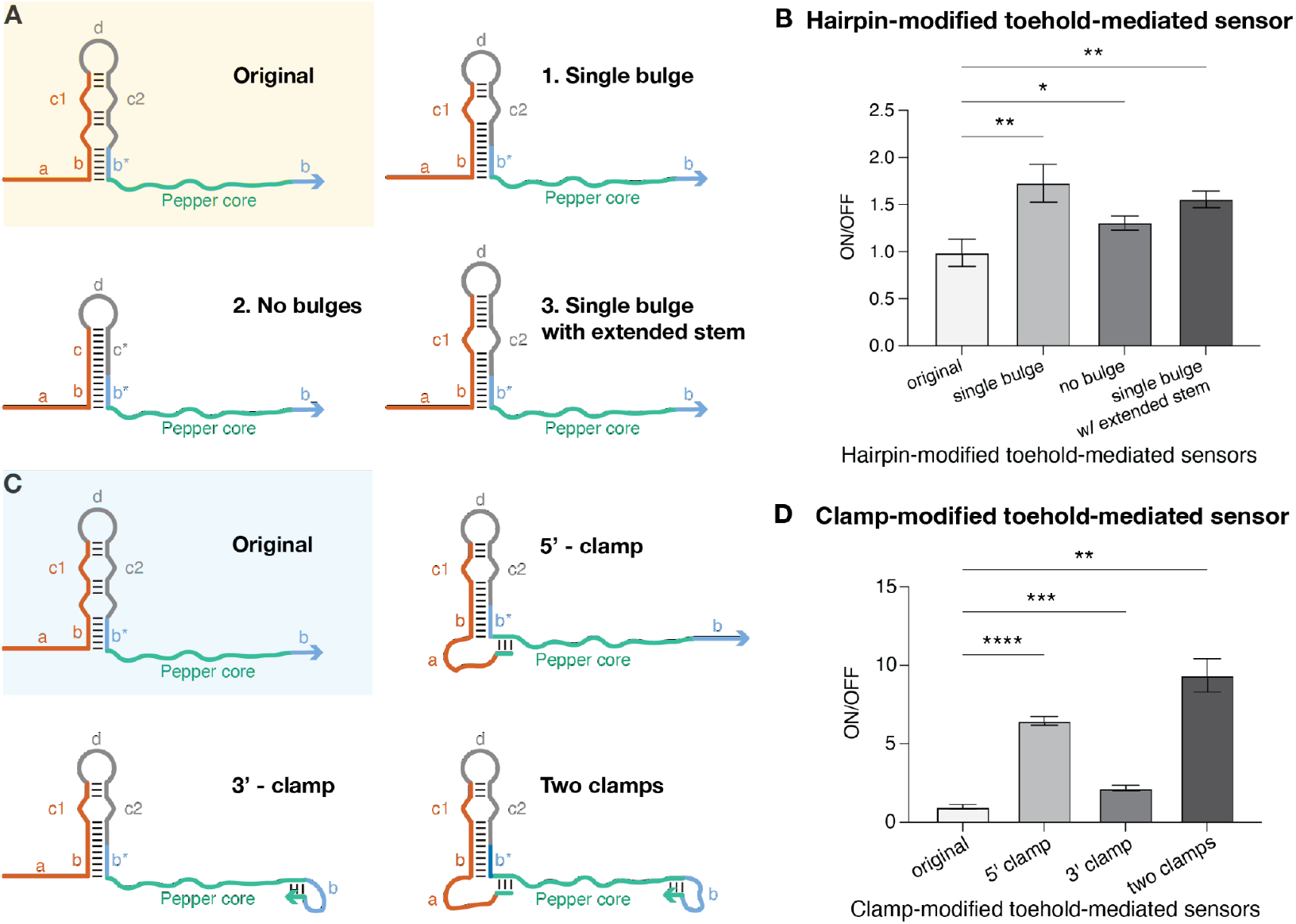
Forward-engineered toehold-mediated Pepper sensors. (A) Schematic of the design modifications made for the forward-engineered sensors. (B) Relative ON/OFF fluorescence ratio obtained for modified toehold-mediated Pepper sensors after two hours of incubation. Error bars represent standard deviations from three technical replicates. (C) Schematic of the design modifications made for the sensors with single bulge. (D) Relative ON/OFF fluorescence ratio obtained for further modified toehold-mediated Pepper sensors after two hours of incubation. Error bars represent standard deviations from three technical replicates. Significance: (****) for p < 0.0001, (***) for p < 0.001, (**) for p < 0.01, (*) for p < 0.1.

We then re-examined the secondary structure predicted by NUPACK of all sensors we evaluated, and we noticed the structure-dependent factors that affect the performance of the sensors. The sensor leakage was decreased when the sequence at the two ends was predicted to interact with the rest of the sequence. We introduced these findings into the sensor design, as shown in Figure 3C, and we referred to these new modifications as clamps. A three-nucleotide sequence complementary to the Pepper core was added upstream of the sensing domain (orange) of the sensor for the 5’ clamp design, whereas a three-nucleotide sequence complementary to the Pepper core was added downstream of the **b** domain (blue) at the 3’ end of the sensor for the 3’ clamp design. When compared with the original sensor #1, the sensor with a single bulge and two clamps showed a significant increase of relative ON/OFF of 948% (Figure 3D). Our results suggest that the improvement in sensor performance achieved through the modification of the sensor stem and the addition of clamps to the sensor design was primarily due to the reduction in signal leakage (Figure S1E), rather than an enhancement in the brightness of the ON signal.

### 2.3 Design of loop-mediated Pepper sensors

Although toehold-mediated Pepper sensors have a wide dynamic range and an ON/OFF ratio exceeding 100-fold (sensors #2 in Fig. 2B and #3 in Fig. 2D), their design has certain limitations regarding the stem sequence of Pepper (**b**/**b**\* domain). In addition, these sensors require a long, single-stranded linear overhang, which is not well-suited for RNA circularization techniques that are commonly used to enhance RNA stability and expression in mammalian cells (35). To overcome these limitations, we developed an alternative design of the Pepper sensor that takes inspiration from loop-initiated RNA activators (LIRA) (33) and builds on earlier RNA sensors based on Broccoli aptamers (29). By placing the target-binding region within the loop of the hairpin structure, the stem sequence of Pepper (**b**/**b**\* domain) in the sensor becomes completely independent of the sequence of the cognate RNA target. Moreover, LIRAs have lower translational leakage compared to toehold-initiated riboregulators (33). Therefore, if they could provide comparable ON signals to the toehold-mediated sensors, we anticipated that the loop-mediated sensors would exhibit higher dynamic range.

Similar to the toehold-mediated sensor, the loop-mediated Pepper sensor includes a hairpin structure at the 5’ end to prevent folding of the downstream Pepper aptamer (Figure 4A). However, we began with a 27-nt long hairpin stem, and a 21-nt long loop, adapting this “first generation” sensor from the LIRA design. In this case, a 24-nt target RNA binds to exposed bases in the loop of the sensor RNA (orange domain), unwinding the hairpin stem and releasing the **b\*** domain. The **b\*** domain in is expected to hybridize with the downstream **b** domain, facilitating the formation of the Pepper aptamer and the activation of the fluorescence. To evaluate this design idea, we generated loop-mediated sensor candidates using a NUPACK script and tested the three sensor-target variants with the lowest ensemble defects. At this stage, the target sequence was varied as part of the sequence optimization program.

**Figure 4.**
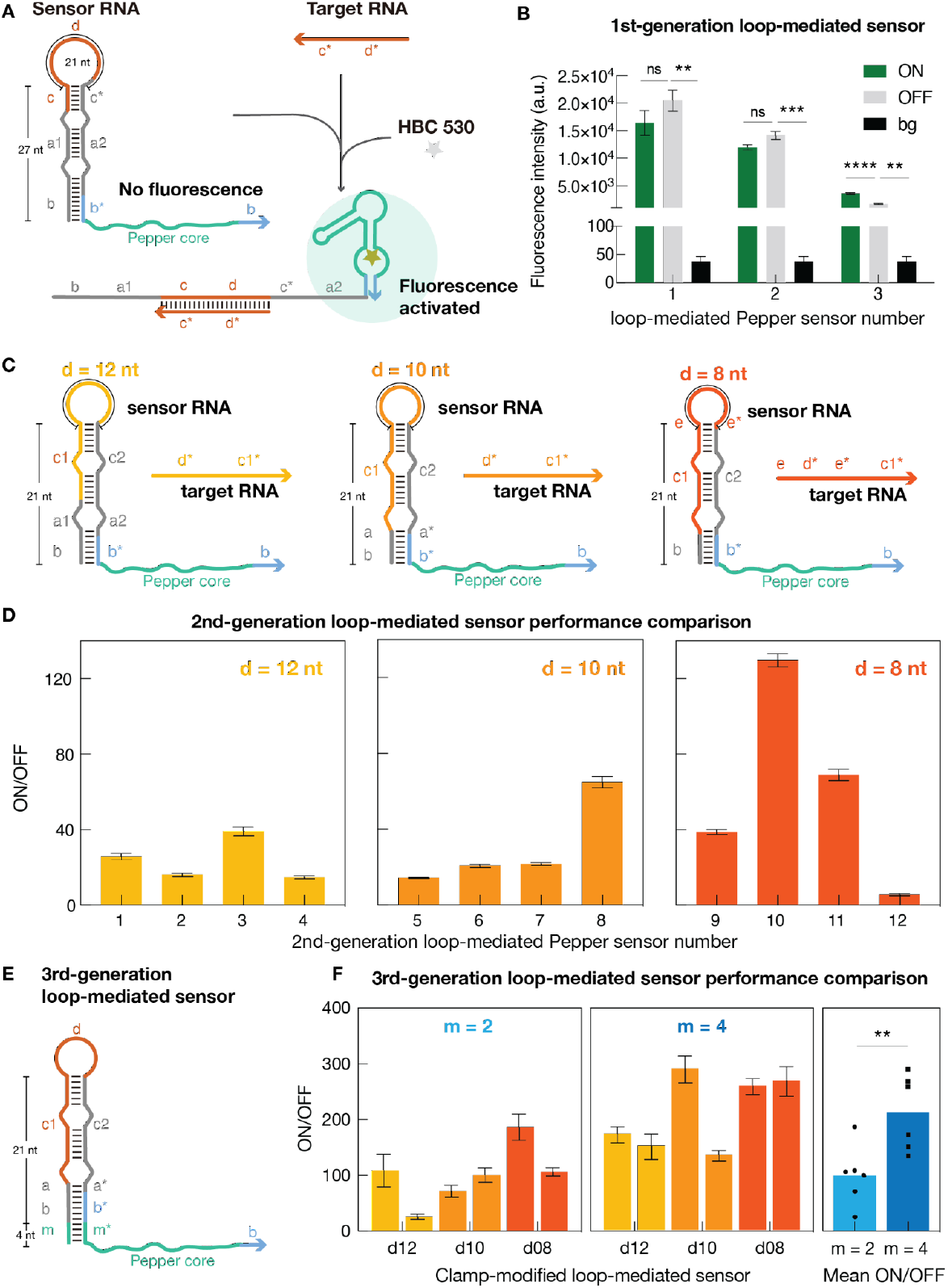
Development of loop-mediated Pepper sensors. (A) Detailed schematic of the loop-mediated Pepper sensor targeting 24-nucleotide target RNA. (B) Best performing loop-mediated Pepper sensors targeting arbitrary RNA targets. Data represent mean fluorescence intensity from plate reader measurements with sensor alone (OFF) and sensor plus 35-nucleotide target RNA (ON). Here “bg” is the fluorescence signal from the HBC530 buffer in the absence of RNA. Error bars represent standard deviations from three technical replicates. (C) Schematic of the second-generation loop-mediated Pepper sensor targeting 24-nucleotide target RNA with various loop sizes. (D) Best performing loop-mediated Pepper sensors targeting shorter RNA targets. Data represent mean fluorescence intensity from plate reader measurements with sensor alone (OFF) and sensor plus 24-nucleotide target RNA (ON). Error bars represent standard deviations from three technical replicates. (E). Schematic of the third-generation loop-mediated Pepper sensor that targets 24-nucleotide RNA. The size of the loop (**d** domain) is 12-, 10- or 8-nt; the length of the clamp (**m** domain) is 4- or 2-nt. (F) The third-generation loop-mediated Pepper sensors and the mean ON/OFF comparison between two clamp designs. Data represent mean fluorescence intensity from plate reader measurements with sensor alone (OFF) and sensor plus 24-nucleotide target RNA (ON). Error bars represent standard deviations from three technical replicates. Significance: (****) for p < 0.0001, (**) for p < 0.01, (ns) for not significant.

As shown in Fig. 4B, each of the initial variants showed significant signal leakage in absence of the target RNA, which in variants #1 and #2 surpassed the fluorescence in the ON state. To overcome this poor performance, we revised the sensor design parameters following steps similar to those taken for the toehold-mediated sensor, obtaining a “second generation” set of sensors. For these sensors while keeping the target sensing region length unchanged at 24-nt, we reduced the length of the hairpin (21 nt, two helical turns), making it comparable to the stem of the toehold-mediated Pepper sensor (Figure 4C). To generate a generalized design strategy for any given targets, we considered three loop sizes, 12, 10, and 8-nt, and for each size we generated and screened four sensor candidates with arbitrary target sequence (the target domain was not conserved, rather it was included in the NUPACK sequence optimization). Figure 4D shows that these adjustments immediately resulted in an increase of the ON/OFF ratio with a significant decrease of the OFF signal nearly 100-fold (Figure S1F). Our results suggested that our first-generation design of the 27-nt stem was not strong enough to hold a 21-nt loop in place and the fluorescent configuration is more favored.

We next hypothesized that the sensor performance could be further improved by limiting the spontaneous interaction of the **b\*** domain with downstream **b** domain, like in the toehold-mediated design. For this purpose, we build a series of third generation sensors that include a clamp (**m/m\*** domain) at the bottom of the hairpin (Figure 4E). We evaluated 2- or 4-nt long **m/m\*** domains that are complementary to the Pepper core sequence. For each case, we tested two candidate sensors against arbitrary target sequences (i.e. the target sequence was optimized with the sensor domains). We observed greatly improved sensor performance compared to first- and second-generation loop-mediated sensors, with ultralow signal leakage detected (Figure 4F and S1F). On average (across arbitrary targets) the sensors with 4-nt clamps had higher ON/OFF ratios compared to the ones with 2-nt clamps.

### 2.4 Pepper sensors for detecting sequence-specific RNA targets

Having confirmed that our design approach for Pepper sensors allows us to improve their ON/OFF ratio for arbitrary targets, we shifted our attention to building sensors for detecting specific, biologically relevant RNA targets. We developed sensors to detect an MS2-repeat sequence and the mRNA of TUBA1b; long non-coding RNAs (lncRNAs) MALAT1 and mouse and human Xist/Xist RNA; and three microRNAs miR302-a, miR-294 and miR124. MS2-repeat sequences (consisting of 24 MS2 aptamer repeats) are widely used as a tag for live cell mRNA imaging in conjunction with fluorescent protein labeled MS2 coat protein (36, 37). The TUBA1b mRNA was selected because of its ubiquitous importance across organisms: α/β-tubulin heterodimers are the basis of the dynamic cytoskeletal polymers – microtubules that are involved in various cellular functions (38). MALAT1 is one of the best characterized lncRNAs, for its association with several types of human cancers (39). Xist is another well-studied lncRNA which induces the transcriptional silencing of genes on one X chromosome in female cells (40). We chose to detect the A-repeat sequence of the Xist RNA which recruits RNA-binding proteins necessary for the induction of X-linked gene silencing (41). The Xist A-repeat region holds 7.5 and 8.5 copies of a conserved 26-nt sequence separated by U-rich linkers for human and mouse, respectively (42). We targeted these conserved domains and the specific sequence used for detection can be found in the Supplementary Information (Pepper DNA template sequence). For each of these cases, we developed different Pepper RNA sensors (toehold and loop-mediated) following the workflow outlined in previous sections (Figure 5A).

**Figure 5.**
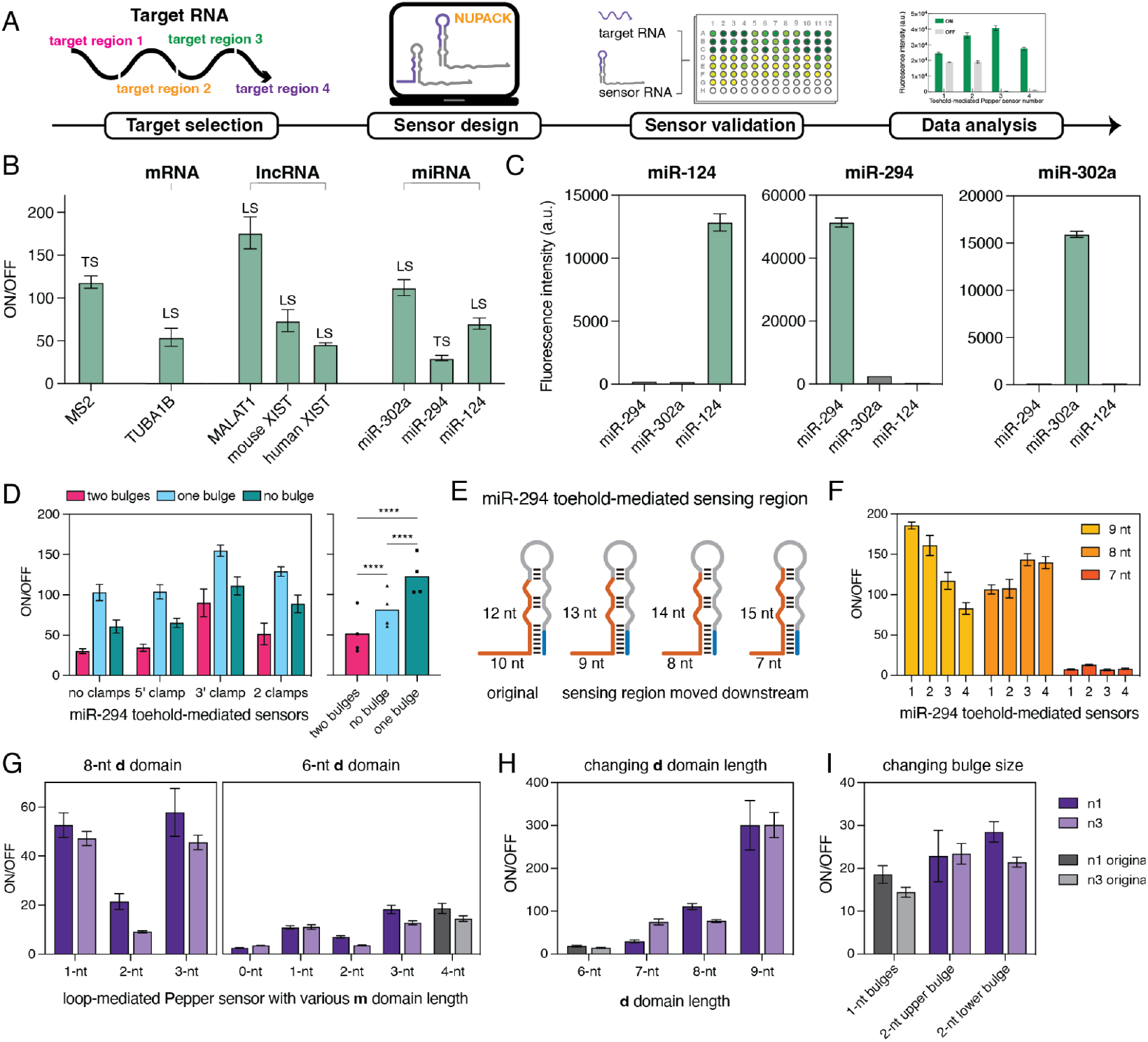
Optimization of Pepper sensors for detection of clinical relevant RNA targets. (A)Illustration of the workflow for developing a Pepper sensor for the detection of a sequence-specific RNA target. (B) The ON/OFF ratios of the best performing sensors for each synthetic RNA target. TS = toehold-mediated sensor; LS = loop-mediated sensor. (C) Specificity test for the three miR sensors measuring the fluorescence in presence of each miR target. (D) The performance of the second-generation toehold-mediated sensors for detecting miR-294. (E) schematic of the modification of the third-generation toehold-mediated sensors for detecting miR-294. (F) The ON/OFF ratios of the third-generation toehold-mediated sensors. (G) The ON/OFF ratios of the loop-mediated sensors with various **m** domain lengths with 8 or 6 nt **d** domain. (H) The ON/OFF ratios of the loop-mediated sensors with **d** domains of 6, 7, 8, or 9 nt (4 nt **m** domain). (I) The ON/OFF ratios of the loop-mediated sensors with a bulge size of 1 or 2 nt (4 nt **m** domain and 6 nt **d** domain). Measurements were taken 2h after mixing in the plate reader. Error bars represent standard deviations from three technical replicates. Significance: (****) for p < 0.0001.

For long target RNAs, such as MS2-repeat, TUBA1B and MALAT, we selected respectively four, two and three target 20- or 24-nt sub-sequences from the full-length product. For shorter target RNAs, such as A-repeats of Xist RNA and the microRNAs, the targeting region is limited so the target sequence is uniquely specified and identical to the full-length product. We validated and screened these sensors in the presence of 1x synthetic (*in vitro* transcribed) RNA targets, and Fig. 5B shows the performance of the best sensors from a pool of three or four original toehold- and 3rd-generation loop-mediated sensors for each RNA target sequence (the performance of the sensors is shown in Fig. S6-9, and sensor sequences can be found in the SI table). Six out of the eight sensors provide ON/OFF ratios over 50-fold upon detecting their specific target, and without any sensor optimization the loop-mediated sensor for MALAT1 had an ON/OFF ratio of ∼176-fold. Moreover, the Pepper sensors for the microRNA showed very high specificity, as they can only be fully switched ON by their cognate RNA target when challenged with non-cognate microRNA targets (Figure 5C).

We next tried to improve the best toehold-mediated Pepper sensor for miR-294, which showed the lowest ON/OFF ratio (30-fold) when compared to other sensors we tested (Figure 5B). We hypothesized that this is due to a combination of a high leakage signal in the absence of target, and low on signal in the presence of target. We found high leakage to affect the ON/OFF ratio of all the toehold-mediated sensors we tested against synthetic miR-294 targets (Figure S2). To reduce the leakage of the toehold-mediated sensors, we added clamps and reduced the number of the bulges in the sensing stem of the sensor. We observed an improvement in performance with a smaller number of bulges and the addition of the 3’ clamp (Figure 5D). Further, we noticed the **b** domain of the sensor contained three consecutive Gs and four Gs in total (5’GGGAAG) which explains why a 3’ clamp was effective. We then moved the sensing region down the sensor by 1-nt at a time (Figure 5E), so the three Gs were moved away from the bottom of the sensing stem. These newly designed sensors had improved performance and exhibited reduced leakage only when the three Gs were flanked by other sequences within the **b** domain (sensors with 9- and 8-nt toehold) as shown in Figure 5F.

To enhance the signal intensity of the loop-mediated miR-294 sensors, we applied the design principles that we previously validated to the loop-mediated sensor N1 and N3: 1) shortening the length of the **m** domain from 4-nt to 3-, 2-, 1- or 0-nt; 2) increasing the length of the **d** domain from 6-to 7-, 8- or 9-nt; and 3) changing the size of bulges within the sensing stem from 1-to 2-nt (Figure 5G-I). The sensors with the **m** domain removed had ∼5-fold increase in ON signal for both N1 and N3, but they also had a ∼37- and ∼20-fold increase in OFF state signal, respectively. Therefore, removing the **m** domain along had an overall negative effect on the system. The sensors with a shortened **m** domain or an enlarged bulge had minimal effect on the brightness of the ON signal, whereas the sensors with longer **d** domain had various degrees of signal enhancement. The best two sensors with 9-nt **d** domain modification displayed ON/OFF ratios of ∼300-fold. We also sought to improve the ON/OFF ratio of the Xist RNA sensors. The conserved sequences within the Xist A-repeat region created a non-linear secondary structure preventing the binding of the target RNA. We applied similar sequence-based modifications to the sensor design (as we did for the miR-294 sensors), and the ON/OFF of the second-generation loop-mediated sensors for the mouse Xist A-repeats improved by ∼17-fold (Fig. S3A-D). Our optimization strategies successfully improved the performance of both the Xist RNA sensors and the miR-294 sensors, demonstrating their potential applicability for any target-specific sensors.

## 3. DISCUSSION

Here we developed two types of Pepper-based fluorogenic RNA sensors that provide RNA detection capabilities *in vitro*. Both toehold- and loop-mediated Pepper sensors exhibit a wide dynamic range. We demonstrated the *in vitro* detection of RNA targets of lengths varying between 20 and 35-nt. We devised approaches to optimize toehold- and loop-mediated sensors’ performances for arbitrary presented target RNA, building upon previous work that established such RNA sensors for diagnostic applications using other aptamers (28–30). Our toehold-mediated sensor design originally included two 2-nt bulges in the hairpin structure to prevent premature transcription termination (43) and increase the thermodynamic stability of target-sensor interactions. However, our results suggest that the ON/OFF ratio of these sensors can be improved by approximately 10-fold through the removal of the bottom bulge in the hairpin structure and the incorporation of two clamps (Fig. 3D). After two rounds of successive improvement, especially with the addition of the clamp at the bottom of the sensor stem, 10 out of 12 of the 3rd generation loop-mediated sensors we tested showed over 100-fold enhancement in fluorescence upon hybridizing with their cognate target RNA. Due to their low background fluorescence in the absence of target RNA, six of these sensors exhibit an ON/OFF ratio exceeding 150-fold. Furthermore, we developed Pepper sensors for detecting various mRNA, lncRNA and microRNA targets with high specificity (Fig. 5C).

The Pepper sensor exhibits a signal-to-background ratio exceeding that of a superquenched DNA molecular beacon (44). The highest performing superquenched molecular beacon, utilizing FAM -3 DABCYLs, achieved a fluorescence signal enhancement of 320-fold through background fluorescence subtraction. Our second-generation loop-mediated Pepper sensors, utilizing a domain **d** of 9-nt, can achieve a fluorescence signal enhancement of approximately 550-fold through background fluorescence subtraction. Additionally, the Pepper sensor is pure RNA-based, eliminating the need for chemical synthesis.

In comparison to previously developed fluorescent light-up RNA-based sensors for detecting microRNA (18, 19, 22, 23), both of our optimized toehold-mediated and loop-mediated sensors demonstrated significantly greater ON/OFF ratios *in vitro*. Because these previously developed sensors have demonstrated utility in live-cell RNA localization or ratiometric imaging applications (11, 12, 16, 18, 19, 21, 22, 24–27), we anticipate that our Pepper sensors could be a valuable tool for RNA detection and tracking *in vivo*. To support this claim, we assessed the sensitivity of the Pepper sensors by conducting target RNA titration experiments using our highest performing sensors. As shown in Fig. S4, our results indicated that all six of the selected sensors at 1 μM were activated by a target RNA concentration as low as 50 nM, which is below the reported concentration of target RNA in the literature (45). Achieving this sensor/target RNA ratio is feasible when expressing the sensor in a medium or high copy number plasmid *in vivo (46)*. Furthermore, we evaluated the robustness of the Pepper sensors in a complex RNA environment by spiking the target RNA into total RNA extracted from HEK 293 cells at various ratios [target]:[background]. We observed no significant change in sensor performance, and the sensors maintained their detection capabilities with up to 50-fold the amount of total RNA (Fig. S5).

Additional studies are needed to verify the specificity of the sensor for detection of RNA in living cells; depending on the applications, discrimination of targets with small mutations may be desirable, or it may be preferable to detect a range of targets with a broad set of mutations. For expression in cells, our sensors will need to be modified to increase their stability, for example by including a three-way junction motif or a tRNA motif that are known to enhance the stability of the ON-state aptamers (47). To optimize sensors for *in vivo* RNA detection, we expect that our *in vitro* design/test/revise cycle will be useful to improve their signal-to-background ratio even when they include additional motifs. Although at the moment other sensors have greater sensitivity than the sensors we demonstrated here (19), our rational design pipeline made it possible for us to build sensors that have a very high signal/background ratio *in vitro*, a parameter that is important in complex samples that could have a high level of background sensing. Our study lays groundwork for the advancement of RNA sensors that exhibit exceptional specificity and sensitivity towards diverse RNA targets. Furthermore, these sensors have the potential to be universally applicable across multiple organisms.

## ACKNOWLEDGEMENTS

This research was supported by the Rose Hill Foundation, by the UCLA Broad Stem Cell Research Center, and by NSF award MCB 2020039 to EF. We thank Alexander Green and Ming Hammond for comments on the manuscript.

## REFERENCES

1. Babendure, J.R., Adams, S.R. and Tsien, R.Y. (2003) Aptamers switch on fluorescence of triphenylmethane dyes. J. Am. Chem. Soc., 125, 14716–14717.

2. Grate, D. and Wilson, C. (1999) Laser-mediated, site-specific inactivation of RNA transcripts. Proc. Natl. Acad. Sci. U. S. A., 96, 6131–6136.

3. Yerramilli, V.S. and Kim, K.H. (2018) Labeling RNAs in Live Cells Using Malachite Green Aptamer Scaffolds as Fluorescent Probes. ACS Synth. Biol., 7, 758–766.

4. Filonov, G.S., Moon, J.D., Svensen, N. and Jaffrey, S.R. (2014) Broccoli: rapid selection of an RNA mimic of green fluorescent protein by fluorescence-based selection and directed evolution. J. Am. Chem. Soc., 136, 16299–16308.

5. Paige, J.S., Wu, K.Y. and Jaffrey, S.R. (2011) RNA mimics of green fluorescent protein. Science, 333, 642–646.

6. Strack, R.L., Disney, M.D. and Jaffrey, S.R. (2013) A superfolding Spinach2 reveals the dynamic nature of trinucleotide repeat-containing RNA. Nat. Methods, 10, 1219–1224.

7. Warner, K.D., Sjekloca, L., Song, W., Filonov, G.S., Jaffrey, S.R. and Ferré-D’Amaré, A.R. (2017) A homodimer interface without base pairs in an RNA mimic of red fluorescent protein. Nat. Chem. Biol., 13, 1195–1201.

8. Dolgosheina, E.V., Jeng, S.C.Y., Panchapakesan, S.S.S., Cojocaru, R., Chen, P.S.K., Wilson, P.D., Hawkins, N., Wiggins, P.A. and Unrau, P.J. (2014) RNA mango aptamer-fluorophore: a bright, high-affinity complex for RNA labeling and tracking. ACS Chem. Biol., 9, 2412–2420.

9. Autour, A., C Y Jeng, S., D Cawte, A., Abdolahzadeh, A., Galli, A., Panchapakesan, S.S.S., Rueda, D., Ryckelynck, M. and Unrau, P.J. (2018) Fluorogenic RNA Mango aptamers for imaging small non-coding RNAs in mammalian cells. Nat. Commun., 9, 656.

10. Chen, X., Zhang, D., Su, N., Bao, B., Xie, X., Zuo, F., Yang, L., Wang, H., Jiang, L., Lin, Q., et al. (2019) Visualizing RNA dynamics in live cells with bright and stable fluorescent RNAs. Nat. Biotechnol., 37, 1287–1293.

11. Alam, K.K., Tawiah, K.D., Lichte, M.F., Porciani, D. and Burke, D.H. (2017) A Fluorescent Split Aptamer for Visualizing RNA-RNA Assembly In Vivo. ACS Synth. Biol., 6, 1710–1721.

12. Wang, Z., Luo, Y., Xie, X., Hu, X., Song, H., Zhao, Y., Shi, J., Wang, L., Glinsky, G., Chen, N., et al. (2018) In Situ Spatial Complementation of Aptamer-Mediated Recognition Enables Live-Cell Imaging of Native RNA Transcripts in Real Time. Angew. Chem. Int. Ed Engl., 57, 972–976.

13. Yu, Q., Ren, K. and You, M. (2021) Genetically encoded RNA nanodevices for cellular imaging and regulation. Nanoscale, 13, 7988–8003.

14. Kolpashchikov, D.M. (2005) Binary malachite green aptamer for fluorescent detection of nucleic acids. J. Am. Chem. Soc., 127, 12442–12443.

15. Kikuchi, N. and Kolpashchikov, D.M. (2017) A universal split spinach aptamer (USSA) for nucleic acid analysis and DNA computation. Chem. Commun., 53, 4977–4980.

16. Karunanayake Mudiyanselage, A.P.K.K., Yu, Q., Leon-Duque, M.A., Zhao, B., Wu, R. and You, M. (2018) Genetically Encoded Catalytic Hairpin Assembly for Sensitive RNA Imaging in Live Cells. J. Am. Chem. Soc., 140, 8739–8745.

17. Zadeh, J.N., Steenberg, C.D., Bois, J.S., Wolfe, B.R., Pierce, M.B., Khan, A.R., Dirks, R.M. and Pierce, N.A. (2011) NUPACK: Analysis and design of nucleic acid systems. J. Comput. Chem., 32, 170–173.

18. Huang, K., Doyle, F., Wurz, Z.E., Tenenbaum, S.A., Hammond, R.K., Caplan, J.L. and Meyers, B.C. (2017) FASTmiR: an RNA-based sensor for in vitro quantification and live-cell localization of small RNAs. Nucleic Acids Res., 45, e130.

19. Ying, Z.-M., Wu, Z., Tu, B., Tan, W. and Jiang, J.-H. (2017) Genetically Encoded Fluorescent RNA Sensor for Ratiometric Imaging of MicroRNA in Living Tumor Cells. J. Am. Chem. Soc., 139, 9779–9782.

20. Bhadra, S. and Ellington, A.D. (2014) A Spinach molecular beacon triggered by strand displacement. RNA, 20, 1183–1194.

21. Kitto, R.Z., Christiansen, K.E. and Hammond, M.C. (2021) RNA-based fluorescent biosensors for live cell detection of bacterial sRNA. Biopolymers, 112, e23394.

22. Dou, C.-X., Liu, C., Ying, Z.-M., Dong, W., Wang, F. and Jiang, J.-H. (2021) Genetically Encoded Dual-Color Light-Up RNA Sensor Enabled Ratiometric Imaging of MicroRNA. Anal. Chem., 93, 2534–2540.

23. Aw, S.S., Tang, M.X., Teo, Y.N. and Cohen, S.M. (2016) A conformation-induced fluorescence method for microRNA detection. Nucleic Acids Res., 44, e92.

24. Ong, W.Q., Citron, Y.R., Sekine, S. and Huang, B. (2017) Live Cell Imaging of Endogenous mRNA Using RNA-Based Fluorescence ‘Turn-On’ Probe. ACS Chem. Biol., 12, 200–205.

25. Sato, S.-I., Watanabe, M., Katsuda, Y., Murata, A., Wang, D.O. and Uesugi, M. (2015) Live-cell imaging of endogenous mRNAs with a small molecule. Angew. Chem. Int. Ed Engl., 54, 1855–1858.

26. Wang, T. and Simmel, F.C. (2023) Switchable fluorescent light-up aptamers based on riboswitch architectures. Angew. Chem. Int. Ed Engl.

27. Wang, Q., Xiao, F., Su, H., Liu, H., Xu, J., Tang, H., Qin, S., Fang, Z., Lu, Z., Wu, J., et al. (2022) Inert Pepper aptamer-mediated endogenous mRNA recognition and imaging in living cells. Nucleic Acids Res., 50, e84.

28. Yan, Z., Tang, A.A., Eshed, A., Ticktin, Z.M., Chaudhary, S., Ma, D., McCutcheon, G., Li, Y., Wu, K., Saha, S., et al. (2023) Rapid and multiplexed nucleic acid detection using programmable aptamer-based RNA switches. medRxiv, 10.1101/2023.06.02.23290873.

29. Tang, A. (2020) RNA Aptamer-Based Systems for Pathogen Detection and Biomolecule Synthesis. Doctoral thesis, Arizona State University, 21–46.

30. Green, A., Ma, D., Tang, A. (2017) Unimolecular aptamer-based sensors for pathogen detection. US Patent Application. PCT/US2017/056960

31. Green, A. (2021) Molecular fuses for real-time, label-free, multiplexed imaging of RNAs in living cells. US Patent. US 11,047,000

32. Green, A.A., Silver, P.A., Collins, J.J. and Yin, P. (2014) Toehold switches: de-novo-designed regulators of gene expression. Cell, 159, 925–939.

33. Ma, D., Li, Y., Wu, K., Yan, Z., Tang, A.A., Chaudhary, S., Ticktin, Z.M., Alcantar-Fernandez, J., Moreno-Camacho, J.L., Campos-Romero, A., et al. (2022) Multi-arm RNA junctions encoding molecular logic unconstrained by input sequence for versatile cell-free diagnostics. Nat Biomed Eng, 6, 298–309.

34. Lam, J.K.W., Chow, M.Y.T., Zhang, Y. and Leung, S.W.S. (2015) siRNA Versus miRNA as Therapeutics for Gene Silencing. Mol. Ther. Nucleic Acids, 4, e252.

35. Litke, J.L. and Jaffrey, S.R. (2019) Highly efficient expression of circular RNA aptamers in cells using autocatalytic transcripts. Nat. Biotechnol., 37, 667–675.

36. Markaki, Y., Chong, J.G., Wang, Y., Jacobson, E.C., Luong, C., Tan, S.Y.X., Jachowicz, J.W., Strehle, M., Maestrini, D., Banerjee, A.K., et al. (2021) Xist nucleates local protein gradients to propagate silencing across the X chromosome. Cell, 184, 6212.

37. Wu, B., Chao, J.A. and Singer, R.H. (2012) Fluorescence fluctuation spectroscopy enables quantitative imaging of single mRNAs in living cells. Biophys. J., 102, 2936–2944.

38. Kim, N.D., Park, E.-S., Kim, Y.H., Moon, S.K., Lee, S.S., Ahn, S.K., Yu, D.-Y., No, K.T. and Kim, K.-H. (2010) Structure-based virtual screening of novel tubulin inhibitors and their characterization as anti-mitotic agents. Bioorg. Med. Chem., 18, 7092–7100.

39. Amodio, N., Raimondi, L., Juli, G., Stamato, M.A., Caracciolo, D., Tagliaferri, P. and Tassone, P. (2018) MALAT1: a druggable long non-coding RNA for targeted anti-cancer approaches. J. Hematol. Oncol., 11, 63.

40. Loda, A. and Heard, E. (2019) Xist RNA in action: Past, present, and future. PLoS Genet., 15, e1008333.

41. Colognori, D., Sunwoo, H., Wang, D., Wang, C.-Y. and Lee, J.T. (2020) Xist Repeats A and B Account for Two Distinct Phases of X Inactivation Establishment. Dev. Cell, 54, 21–32.e5.

42. Pintacuda, G., Young, A.N. and Cerase, A. (2017) Function by Structure: Spotlights on Xist Long Non-coding RNA. Front Mol Biosci, 4, 90.

43. Lesnik, E.A., Sampath, R., Levene, H.B., Henderson, T.J., McNeil, J.A. and Ecker, D.J. (2001) Prediction of rho-independent transcriptional terminators in Escherichia coli. Nucleic Acids Res., 29, 3583–3594.

44. Yang, C.J., Lin, H. and Tan, W. (2005) Molecular assembly of superquenchers in signaling molecular interactions. J. Am. Chem. Soc., 127, 12772–12773.

45. Bosson, A.D., Zamudio, J.R. and Sharp, P.A. (2014) Endogenous miRNA and target concentrations determine susceptibility to potential ceRNA competition. Mol. Cell, 56, 347– 359.

46. Phillips, G.J., Park, S.K. and Huber, D. (2000) High copy number plasmids compatible with commonly used cloning vectors. Biotechniques, 28, 400–2, 404, 406 passim.

47. Filonov, G.S., Kam, C.W., Song, W. and Jaffrey, S.R. (2015) In-gel imaging of RNA processing using broccoli reveals optimal aptamer expression strategies. Chem. Biol., 22, 649–660.

