## Supplemental Files for "Optimization of RNA Pepper sensors for the detection of arbitrary RNA targets"

### Sequence design

The RNA sequences for the Pepper sensors were designed using NUPACK. The three G's were added at the 5' end of each RNA strand for maximizing the efficiency of the *in vitro* transcription reaction. The secondary structure of the target strand, the sensor strand and the target-sensor complex were specified in DU+ notation. The design scripts listed below were used to generate the toehold-mediated sensors and loop-mediated sensors for detecting arbitrary RNA targets. The target sequence was defined in the scripts when designing sensors for detecting the sequence-specific RNA targets.

------------------------------------------------------------------------------------------------------------------

```
#NUPACK script for the 35-nt toehold-mediated Pepper sensors.

material = rna1999
temperature = 37
trials = 10
structure target = U3 U35 U3 U47
structure sensor = U3 U15 D12 (U1 D3 (U1 D3 U8 U1) U1) U43 U6 U3 U47
structure target_sensor = U3 D35 (U3 U47 + U3) U8 U14 D6 U43 U3 U47

domain T7_term_min = UAGCAUAACCCCUUGGGGCCUCUAAACGGGUCUUGAGGGGUUUUUUG
domain apt_sta_core = ACGGUCGGGUCCAGAUAUUCGUAUCUGUCGAGUAGAGUGUGGG
domain apt_rot_core = UCGAGUAGAGUGUGGGCUCAGAUUCGUCUGAGACGGUCGGGUC

domain threeGs = GGG
domain target_linker = NNN
domain sensor_linker = NNN
domain a = N15
domain b = N6
domain c1 = N14
domain d = N8
domain c2 = N14

target.seq = threeGs c1* b* a* target_linker T7_term_min
sensor.seq = threeGs a b c1 d c2 b* apt_rot_core b sensor_linker T7_term_min
target_sensor.seq = threeGs c1* b* a* target_linker T7_term_min threeGs a b c1 d c2 b* apt_rot_core b
sensor_linker T7_term_min

prevent = AAAA, CCCC, GGGG, UUUU, KKKKKK, MMMMMM, RRRRRR, SSSSSS, WWWWWW,
YYYYYY
```

------------------------------------------------------------------------------------------------------------------

```
#NUPACK script for the loop-mediated Pepper sensors with a 10-nt loop and a 4-nt stem clamp.

material = rna
temperature = 37.0
trials = 10

structure target = U3 U24 U3 U47
structure sensor = U3 D11 (U1 D6 (U1 D6 U8 U1) U1) U37 U6 U3 U47
structure target_sensor = U3 D24 (U3 U47 + U3 U10) U14 D6 U41 U3 U47

domain target_linker = NNN
domain sensor_linker = NNN

domain b = N6
domain c1 = N14
domain c2 = N14
domain d = N8
domain e = N
domain m = N4

domain T7_term_min = UAGCAUAACCCCUUGGGGCCUCUAAACGGGUCUUGAGGGGUUUUUUG
domain apt_rot_core = ACUGGCGCCGCCUGCUUCGGCAGGCCAAUCGUGGCGUGUCG
domain threeGs = GGG

target.seq = threeGs e d* e* c1* target_linker T7_term_min
sensor.seq = threeGs m b c1 e d e* c2 b* apt_rot_core b sensor_linker T7_term_min
target_sensor.seq = threeGs e d* e* c1* target_linker T7_term_min threeGs m b c1 e d e* c2 b*
apt_rot_core b sensor_linker T7_term_min

prevent = AAAA, CCCC, GGGG, UUUU, KKKKKK, MMMMMM, RRRRRR, SSSSSS, WWWWWW,
YYYYYY
```
--------------------------------------------------------------------------------------------------------------------

### Pepper aptamer variations

For the better incorporation of the Pepper into the sensor system, we tested three different versions of the Pepper as illustrated in Fig.S1A. The standard version was the original Pepper aptamer, and the rotated versions were the circularized permutations of the Pepper. We truncated the original stem sequence for the rotated version 1 and kept the stem sequence for the rotated version 2. We replaced the stabilizing stem sequence with 8 scrambled sequences (N1-N8) generated with NUPACK. The fluorescence intensities were measured with 1 μM of the aptamer in 2 μM of HBC 530 buffer (Fig. S1B). The fluorescence intensities of both standard and rotated v2 Pepper were approximately 100-fold than those of the rotated v1 Pepper. Moreover, they were not affected by the scrambled stem sequence replacements. We used standard and rotated v2 Pepper for the sensor designs in the later experiments, and referred to the rotated v2 Pepper as the rotated Pepper. We designed the sensors for detecting 35-nt targets with the incorporation of the standard Pepper.  The four sensors we tested (sta-arb-A–D) all had very high leakage (Fig. S1C), whereas the four sensors with rotated Pepper with the exact same design had very low leakage without any optimization suggesting the rotated Pepper was a better fit for sensor incorporation.

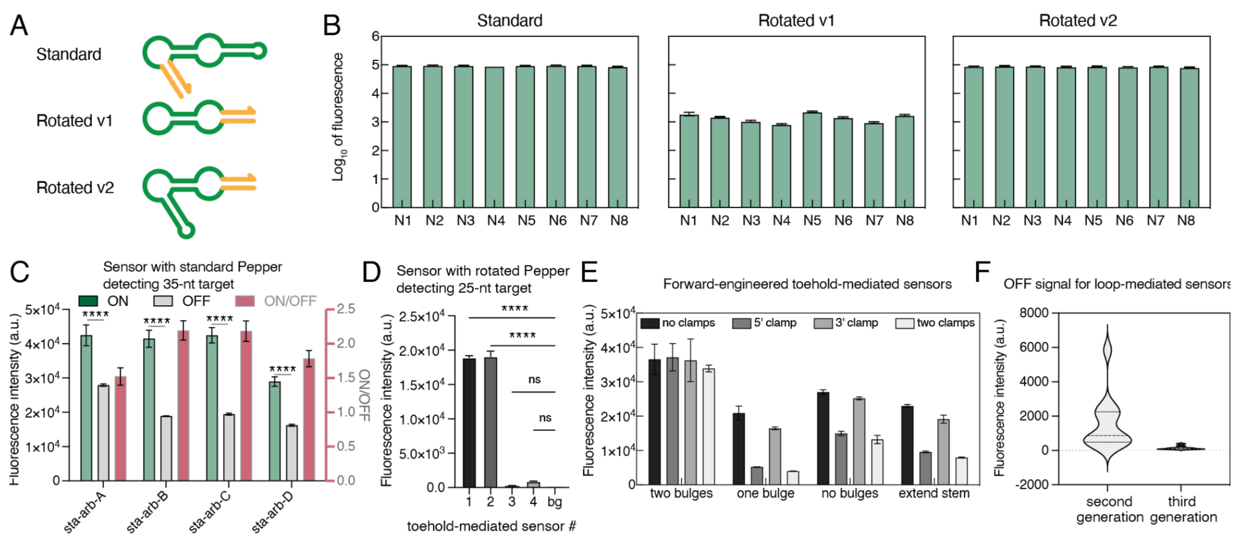


*Figure S1. Pepper variations and the Pepper sensor for arbitrary targets. (A) Secondary structure for the three Pepper variations: standard Pepper, rotated Pepper version 1, and rotated Pepper version 2. The Pepper core sequence is highlighted in green, and the stabilizing stem is in yellow. (B) The fluorescence intensity of the Pepper aptamer with scrambled stem sequence. (C) The performance of the toehold-mediated sensor with standard Pepper for detecting 35-nt target RNA. Data represent mean fluorescence intensity from plate reader measurements with sensor alone (OFF) and sensor plus the  35-nt target RNA (ON). (D) The OFF signal intensities of the toehold-mediated sensor for detecting a 25-nt RNA target employing the rotated Pepper aptamer (v2). The background fluorescence intensity of the HBC350 buffer is listed as bg (background). (E) The OFF signal intensities of the forward-engineered toehold-mediated sensors. (F) The OFF signal intensities of the second and third generation loop-mediated sensors. The adjusted p value was calculated with Šídák's multiple comparisons test. We used the adjusted significance level to control the overall Type I error rate.*

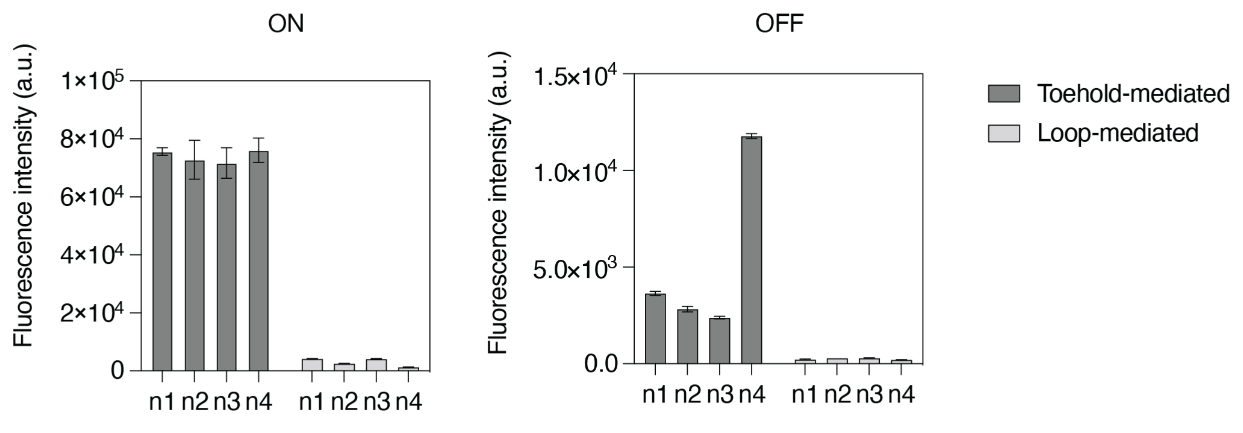


*Figure S2. ON and OFF fluorescence signal of the toehold- and loop-mediated Pepper sensors for detecting miR294. Error bars represent standard deviations from three technical replicates.*

**Optimization of the Xist A-repeat sensors**

Sequence-dependent sensor design optimization was performed for the Xist A-repeat sensors. We selected a 24-nt sequence from the third human A-repeat and a 25-nt sequence from the mouse fourth mouse A-repeat for the Pepper sensor design. The first generation loop-mediated Pepper sensors for both mouse and human A-repeats had an average ON/OFF around 5-fold (Fig. S3A and B). The sensors were not fully switched on because stem-loop structures were formed in the loop region (**d** domain) preventing the invasion of the target RNA strand at the loop. To overcome this sequence constraint, we implemented two approaches to release the unintentional stem-loop structure at the loop: 1) increasing the size of the loop while keeping the target RNA binding to the entire loop; and 2) increasing the size of the loop by the addition of nucleotides downstream of the target binding region of the sensor. The first approach (d18s20, d16s20 and d14s20) changing to loop size to 18-, 16- and 14-nt didn't improve the performance of the mouse A-repeat sensors, and only enhanced the performance of the human A-repeat sensors by approximately two-folds (Fig. S3C and D). One design modification (d10n4) from the second approach (d10n6, d10n5 and d10n4) changing the loop size to 14-nt while keeping the 10-nt target-bind region in the loop and 14-nt target-bind region in the stem, had the most significant improvement to the sensor performance (Fig. S3C and D). The introduction of the random sequence to the loop region in the second approach had better reduced the sequence-dependent structural constraints at the loop. Furthermore, the second approach allowed the target RNA to bind further down to the stem which helped the unwinding of the sensor stem.

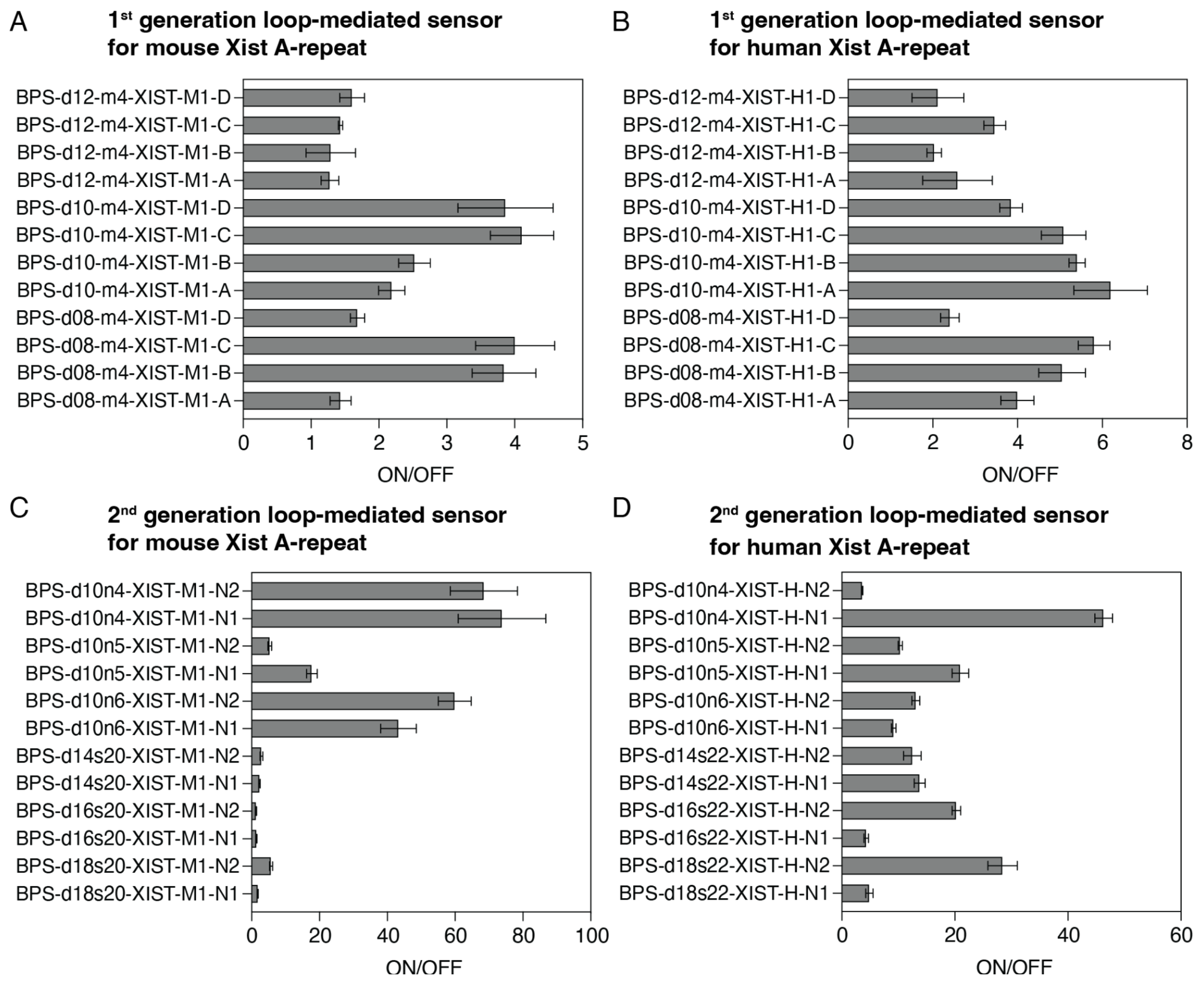


*Figure S3. Performance of the loop-mediated Pepper sensors for detecting Xist A-repeat sequences. (A) 1st generation loop-mediated Pepper sensors for the detection of mouse Xist A-repeat sequences. (B) 1st generation loop-mediated Pepper sensors for the detection of human Xist A-repeat sequences. (C) 2nd generation loop-mediated Pepper sensors for the detection of mouse Xist A-repeat sequences. (D) 2nd generation loop-mediated Pepper sensors for the detection of human Xist A-repeat sequences. Data represent mean fluorescence intensity from plate reader measurements with sensor alone (OFF) and* sensor plus the synthetic Xist A-repeat target RNA (ON). Error bars represent standard deviations from three technical replicates.

### Assessing Pepper sensor sensitivity

We selected six best performing sensors for target RNA titration experiments and found all sensors were activated with as little as 50 nM of synthetic target RNA (Fig.S4 A-F).

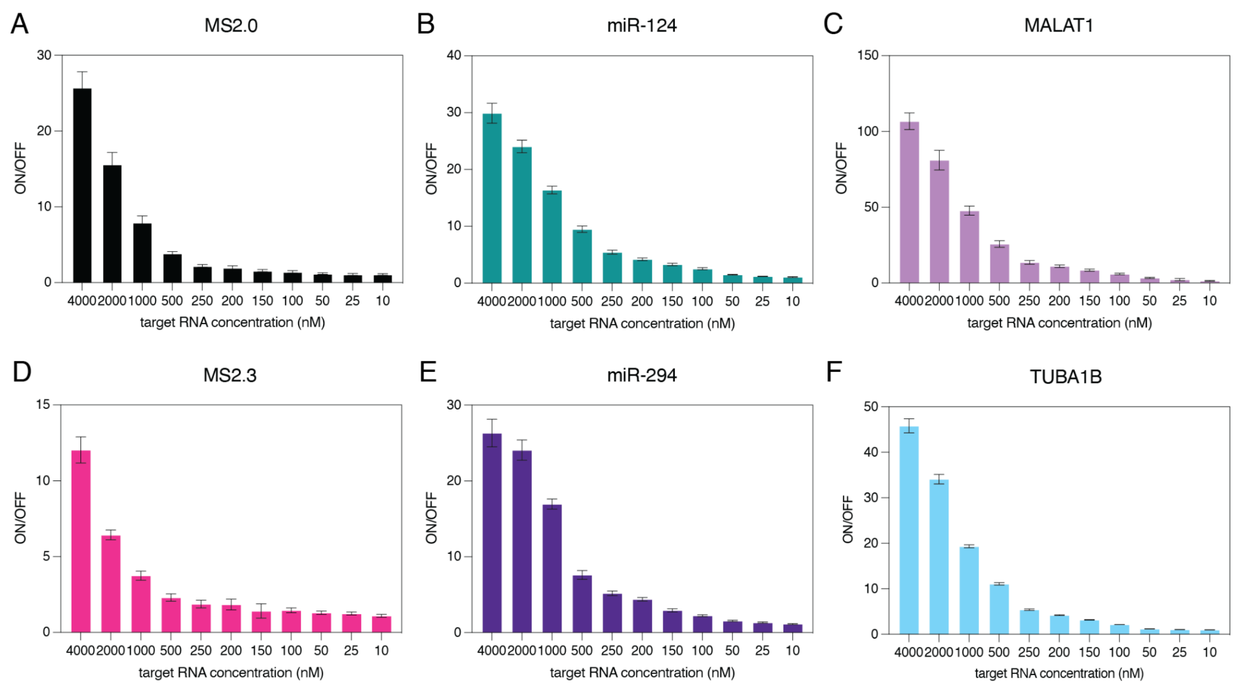


*Figure S4. Detection limit measurements for synthetic target RNAs with 1 μM of sensor RNA. (A) The limit of detection for the MS2.0 sensor. (B) The limit of detection for the miR-124 sensor. (C) The limit of detection for the MALAT1 sensor. (D) The limit of detection for the MS2.3 sensor. (E) The limit of detection for the miR-294 sensor. (F) The limit of detection for the TUBA1B sensor. Data represent mean fluorescence intensity from plate reader measurements with sensor alone (OFF) and sensor plus the cognate synthetic target RNA (ON). Error bars represent standard deviations from three technical replicates.*

### Effects of the background RNA

Synthetic target RNAs were spiked into total RNA extracted from HEK293 cells to evaluate the effect of the background RNA on the Pepper sensor performance. We found no significant differences in ON/OFF even when 50x total RNA was added to 50 nM of target RNA across all four sensors (Fig.S5A-D).

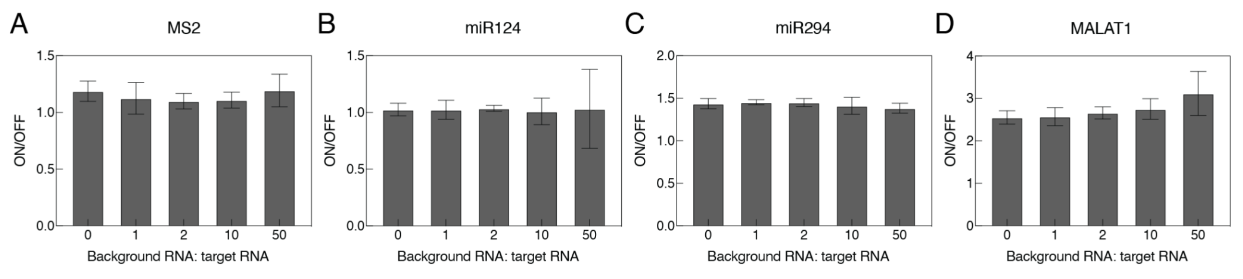


*Figure S5. The effects of the background RNA on the Pepper sensor performance. Error bars represent*

*standard deviations from three technical replicates.*

### Performance of the sensors for detecting sequence-specific RNA targets

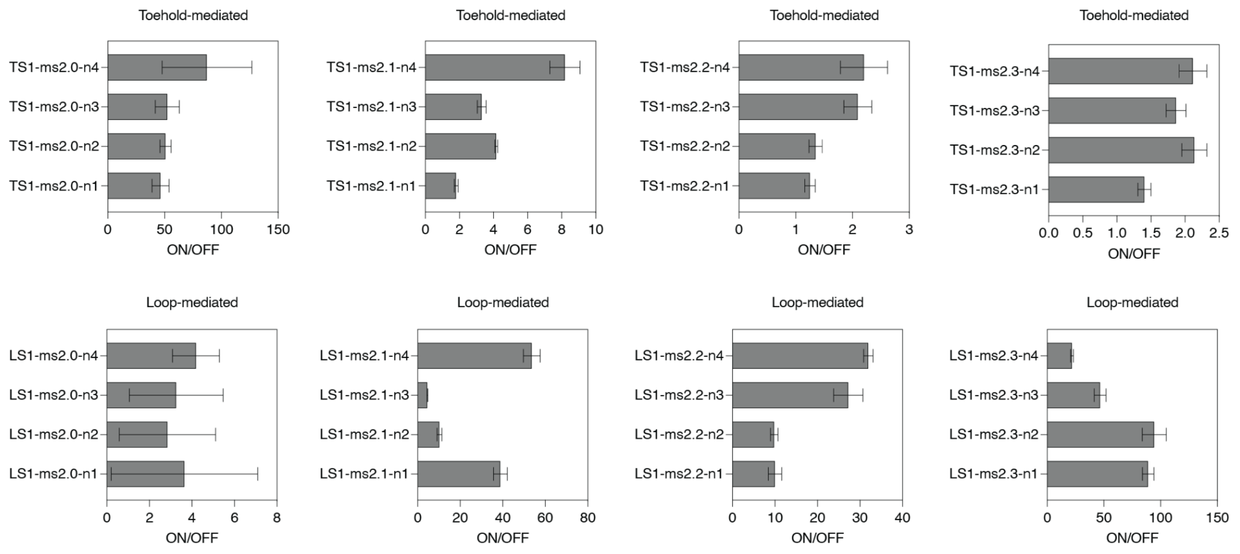


*Figure S6. The Performance of the toehold- and loop-mediated Pepper sensors for detecting four different targeting regions on the MS2 repeat. Error bars represent standard deviations from three technical replicates.*

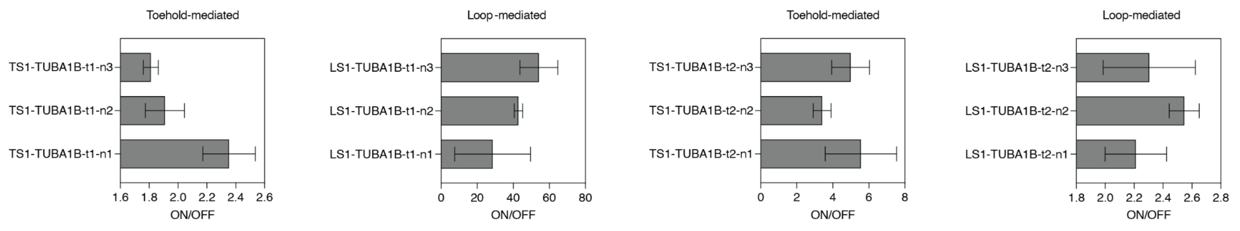


*Figure S7. The Performance of the toehold- and loop-mediated Pepper sensors for detecting two different targeting regions of the mRNA TUBA1B. Error bars represent standard deviations from three technical replicates.*

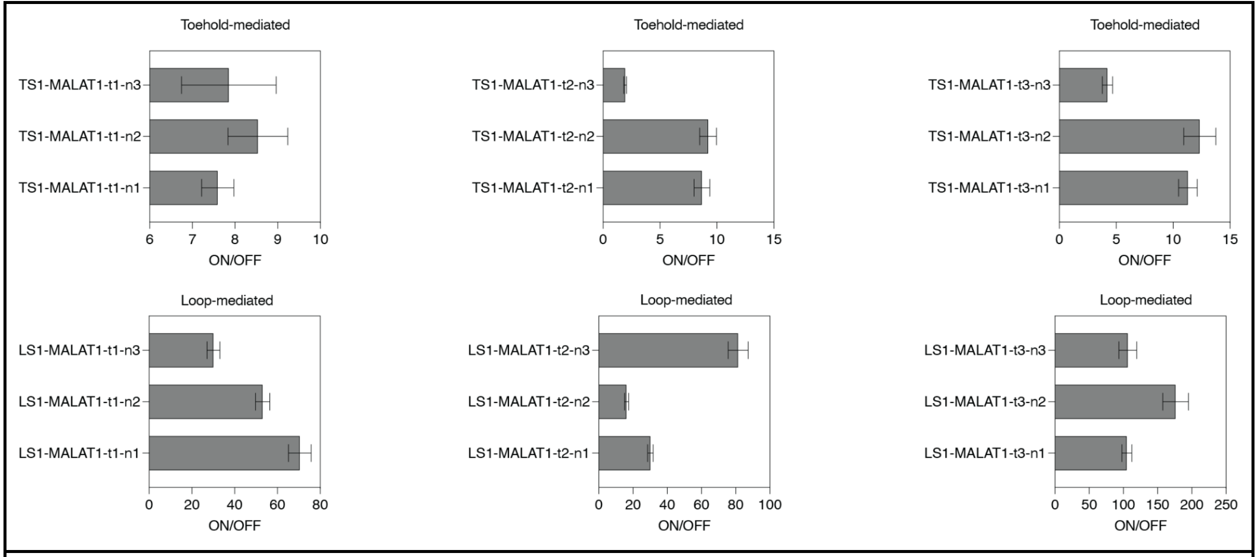


*Figure S8. The Performance of the toehold- and loop-mediated Pepper sensors for detecting three different targeting regions of the mRNA MALAT1. Error bars represent standard deviations from three technical replicates.*

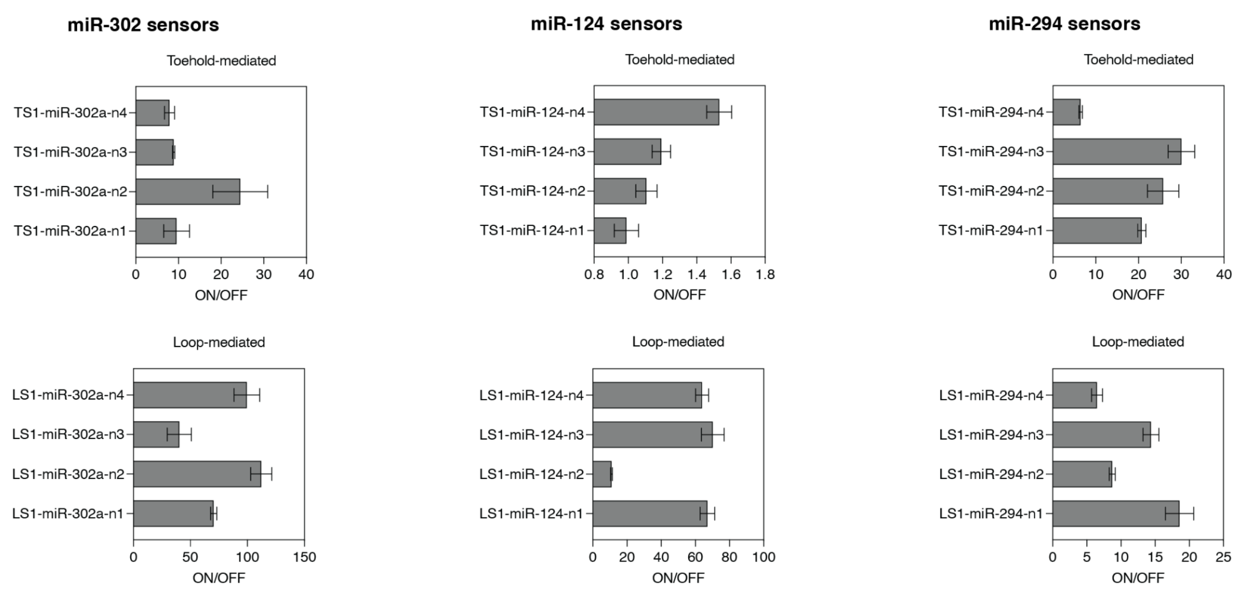


*Figure S9. The Performance of the toehold- and loop-mediated Pepper sensors for detecting miR-302a, miR-124 and miR-294. Error bars represent standard deviations from three technical replicates.*

*Table S1. Overview of the performance of previously developed FLAP-based devices.*

| Reference | Cell type | Fold change (*in vitro*) | Fold change (*in vivo*) | Notes |
|---|---|---|---|---|

| | | | | |
|---|---|---|---|---|
| 11. Alam,K.K., Tawiah,K.D., Lichte,M.F., Porciani,D. and Burke,D.H. (2017) A Fluorescent Split Aptamer for Visualizing RNA-RNA Assembly In Vivo. *ACS Synth. Biol.*, **6**, 1710–1721. | E. Coli | 186 | 6 | Technically not a sensor<br>3WJdB-T insert provided~ 25 fold change *in vivo* relative to pUC19 transformants alone |
| 12. Wang,Z., Luo,Y., Xie,X., Hu,X., Song,H., Zhao,Y., Shi,J., Wang,L., Glinsky,G., Chen,N., *et al.* (2018) In Situ Spatial Complementation of Aptamer-Mediated Recognition Enables Live-Cell Imaging of Native RNA Transcripts in Real Time. *Angew. Chem. Int. Ed Engl.*, **57**, 972–976. | **Mammalian cells:** HeLa cells, HuMSC | ~8 | n/a | |
| 14.Kolpashchikov,D.M. (2005) Binary malachite green aptamer for fluorescent detection of nucleic acids. *J. Am. Chem. Soc.*, **127**, 12442–12443. | n/a | 20 | n/a | |
| 15. Kikuchi,N. and Kolpashchikov,D.M. (2017) A universal split spinach aptamer (USSA) for nucleic acid analysis and DNA computation. *Chem. Commun.* , **53**, 4977–4980. | n/a | 74 | n/a | |
| 16. Karunanayake Mudiyanselage,A.P.K.K., Yu,Q., Leon-Duque,M.A., Zhao,B., Wu,R. and You,M. (2018) Genetically Encoded Catalytic Hairpin Assembly for Sensitive RNA Imaging in Live Cells. *J. Am. Chem. Soc.*, **140**, 8739–8745. | E. coli | 3 | n/a | |

| | | | | |
|---|---|---|---|---|
| 18. Huang,K., Doyle,F., Wurz,Z.E., Tenenbaum,S.A., Hammond,R.K., Caplan,J.L. and Meyers,B.C. (2017) FASTmiR: an RNA-based sensor for in vitro quantification and live-cell localization of small RNAs. *Nucleic Acids Res.*, **45**, e130. | **Mammalian cells:** Huh7 cells, HEK293 or HEK293T | ~10 | | calculated from Figure 3A (background fluorescence subtracted) |
| 19. Ying,Z.-M., Wu,Z., Tu,B., Tan,W. and Jiang,J.-H. (2017) Genetically Encoded Fluorescent RNA Sensor for Ratiometric Imaging of MicroRNA in Living Tumor Cells. *J. Am. Chem. Soc.*, **139**, 9779–9782. | **Mammalian cells:** Hela, MCF-7, L-02 | ~8 | | |
| 20. Bhadra,S. and Ellington,A.D. (2014) A Spinach molecular beacon triggered by strand displacement. *RNA*, **20**, 1183–1194. | n/a | 70 | n/a | **70-fold** increment over the fluorescence of DFHBI alone (with the sensor RNA, presumably) |
| 21.Kitto,R.Z.,Christiansen, K.E.and Hammond,M.C. (2021) RNA-based fluorescent biosensors for live cell detection of bacterial sRNA. *Biopolymers*, **112**, e23394. | E. Coli | ~ 7.4 | ~2x | |
| 22. Dou,C.-X., Liu,C., Ying,Z.-M., Dong,W., Wang,F. and Jiang,J.-H. (2021) Genetically Encoded Dual-Color Light-Up RNA Sensor Enabled Ratiometric Imaging of MicroRNA. *Anal. Chem.*, **93**, 2534–2540. | **Mammalian cells:** Hela, HepG2, L-02 | ~5 | | |
| 23. Aw,S.S., Tang,M.X., | n/a | Ranging from | | Table 1. |

| | | | | |
|---|---|---|---|---|
| Teo,Y.N. and Cohen,S.M. (2016) A conformation-induced fluorescence method for microRNA detection. *Nucleic Acids Res.*, 44, e92. | | 3 to 118 | | |
| 24. Ong,W.Q., Citron,Y.R., Sekine,S. and Huang,B. (2017) Live Cell Imaging of Endogenous mRNA Using RNA-Based Fluorescence ‘Turn-On’ Probe. *ACS Chem. Biol.*, 12, 200–205. | E. Coli | n/a | | |
| 25.Sato,S.-I., Watanabe,M., Katsuda,Y., Murata,A., Wang,D.O. and Uesugi,M. (2015) Live-cell imaging of endogenous mRNAs with a small molecule. *Angew. Chem. Int. Ed Engl.*, 54, 1855–1858. | Mammalian cells: Hela | 5 | | |
| 26.Wang,T. and Simmel,F.C. (2023) Switchable fluorescent light-up aptamers based on riboswitch architectures. *Angew. Chem. Int. Ed Engl.* | E. coli | ~200 (Figure S4a) | ~4 | RNA target and sensor were co-transcribed and the background signal was subtracted (*in vitro* experiment) |
| 27. Wang,Q., Xiao,F., Su,H., Liu,H., Xu,J., Tang,H., Qin,S., Fang,Z., Lu,Z., Wu,J., *et al.* (2022) Inert Pepper aptamer-mediated endogenous mRNA recognition and imaging in living cells. *Nucleic Acids Res.*, 50, e84. | Mammalian cells: Hela, | ~10 | Less than 5-fold | Figure 4D, Figure 6B and E |
| 28. Yan,Z., Tang,A.A., Eshed,A., Ticktin,Z.M., Chaudhary,S., Ma,D., | n/a | ~260 | | Toehold-mediated |

| | | | | |
|---|---|---|---|---|
| McCutcheon,G., Li,Y., Wu,K., Saha,S., *et al.* (2023) Rapid and multiplexed nucleic acid detection using programmable aptamer-based RNA switches. *medRxiv*, 10.1101/2023.06.02.23290873. | | | | |
| 29. Tang,A. (2020) RNA Aptamer-Based Systems for Pathogen Detection and Biomolecule Synthesis. *Doctoral thesis*, Arizona State University, 21-46. | n/a | ~15 | | Loop-mediated |
